# Lack of apoptosis leads to cellular senescence and tumorigenesis in *Drosophila* epithelial cells

**DOI:** 10.1101/2023.05.08.539867

**Authors:** Juan Manuel Garcia-Arias, Noelia Pinal, Sara Cristóbal Vargas, Carlos Estella, Ginés Morata

## Abstract

Programmed cell death (apoptosis) is a homeostasis program of animal tissues designed to remove cells that are unwanted or are damaged by physiological insults. To assess the functional role of apoptosis we have studied the consequences of subjecting *Drosophila* epithelial cells defective in apoptosis to stress or genetic perturbations that normally cause massive cell death. We find that many of those cells acquire persistent activity of the JNK pathway, which drives them into senescent status, characterized by arrest of cell division, cell hypertrophy, Senescent Associated ß-gal activity (SA-ß-gal), ROS production, Senescent Associated Secretory Phenotype (SASP) and migratory behaviour. We have identified two classes of senescent cells in the wing disc: 1) those that localize to the appendage part of the disc, express the *upd*, *wg* and *dpp* signalling genes and generate tumour overgrowths, and 2) those located in the thoracic region do not express *wg* and *dpp* nor they induce tumour overgrowths. Whether to become tumorigenic or non-tumorigenic depends on the original identity of the cell prior to the transformation. We also find that the *p53* gene contributes to senescence by enhancing the activity of JNK.

## INTRODUCTION

Programmed cell death (apoptosis) is a response mechanism to eliminate cells that are unwanted during development [1–3] or are damaged by different stressors such as Ionizing Radiation (IR), heat shock (HS) or drug treatments [4]. The key molecules involved are cysteine proteases (caspases) that dismantle protein substrates and cause death of the affected cells. Caspases are present in all animal cells as inactive zymogens, which may become activated by various factors [5, 6]. The role of caspases as apoptosis executors is conserved in Metazoans [7], although the mechanisms of caspase activation vary between species.

Lack of apoptosis is known to have deleterious effects; it predisposes to the formation of tumours and also causes developmental anomalies [8]. It has been proposed that one of the hallmarks of cancer cells is that they avoid apoptosis [9], thus emphasizing the anti-tumour function of apoptosis. Work in *Drosophila* by us and others [10–13] has also shown that compromising apoptosis often results in tumorous overgrowths.

Because of its sophisticated genetics and readiness for experimental *in vivo* manipulation, *Drosophila* is a convenient system to analyse the consequences of the lack of apoptosis. The apoptosis program is well characterized, especially the response to IR (reviewed in [14, 15]). An outline of the program is shown in Fig. 1. After IR a number of upstream factors become active, which in turn lead to the activation of p53 and of the Jun N-Terminal Kinase (JNK) pathway. These two factors are the major contributors to IR-induced apoptosis; they can elicit apoptosis response independently [16–19], but also stimulate each other [20], what reinforces the apoptotic response. The cellular/genetic events initiated by p53 and JNK to cause cell death are well known: induction of pro-apoptotic genes like *head involution defective (hid)* or *reaper (rpr),* which in turn suppress the activity of the *Drosophila* inhibitor of apoptosis1 (Diap1) protein, thus allowing activation of the apical caspase Dronc and subsequently of the effector caspases Drice and Dcp1 (Fig. 1)(reviewed in [15]).

**Figure 1.**
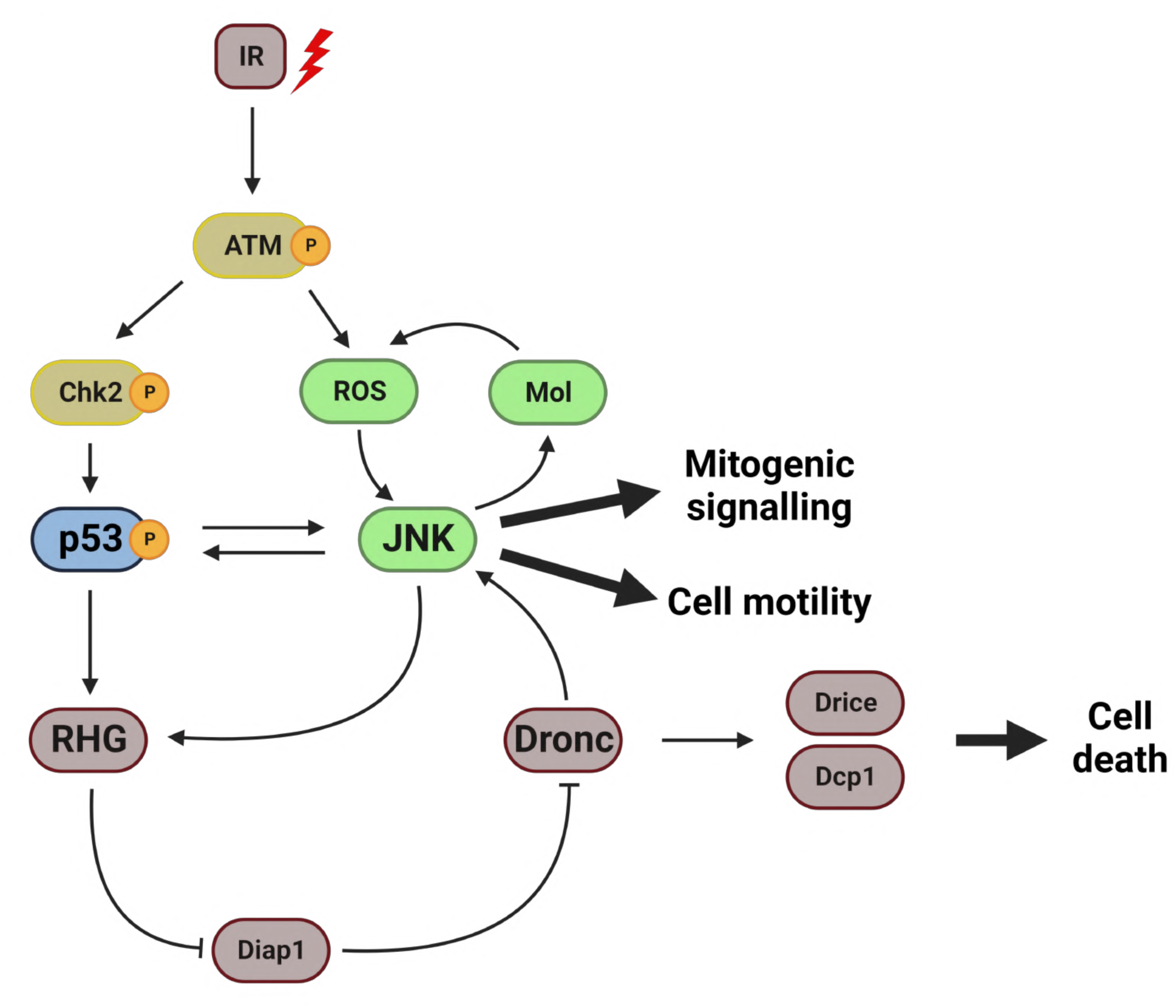
Apoptosis and the Jun N-Terminal Kinase (JNK) pathway in *Drosophila*. Simplified view of the chain of events triggered by the irradiation (IR) of *Drosophila* imaginal cells. The Ataxia telangiectasia mutated (ATM, Tefu in *Drosophila*) protein is activated by IR and in turn causes production of Reactive Oxygen Species (ROS) and induction of Checkpoint kinase 2 (Chk2). The p53 protein and the Jun N-terminal pathway are activated subsequently and mutually stimulate each other. This results in the activation of the pro-apoptotic genes, *reaper, hid* and *grim* (RHG in the Fig.). The function of the latter degrades the product of the *Drosophila inhibitor of apoptosis 1 (diap1)* gene, which allows activation of the apical caspase Dronc and subsequently of the effector caspases Drice and Dcp1, which in turn dismantle protein substrates and cause cell death. Thus, both p53 and JNK exert an autocrine cell killing function. In addition, cells expressing JNK has two other properties: 1) an endogenous activity that stimulates their own movement, a cell motility function, and 2) a secretory function, generating mitogenic signals that promote the proliferation of surrounding cells. In normal circumstances the cell killing function prevents the expression of cell motility and mitogenic signalling, but these two functions can be studied when the apoptotic function is impeded by suppression of the RHG genes or the loss of Dronc activity.

However, and importantly, JNK also has other non-apoptotic roles (Fig. 1, see review in [14]); it has a paracrine function that stimulates the proliferation of cells close to those expressing JNK. This pro-proliferation activity is likely due to secreted growth factors, like Decapentaplegic (Dpp) and Wingless (Wg), known targets of JNK and responsible for the overgrowths caused by “undead” cells, in which cell death is blocked by the effector caspases inhibitor P35 [11–13]. This paracrine function also plays a role in regeneration processes in which dying JNK-expressing cells stimulate the growth of surrounding tissue [21].

Another property associated with JNK activity is that it promotes cell migration, both in vertebrates [22, 23] and in *Drosophila* [13, 24, 25], which may lead to JNK-expressing cells invading neighbour compartments.

Thus, there are at least three distinct functions of JNK that have been characterized when active in imaginal cells: two endogenous (autocrine) activities - pro-apoptosis and pro-migration- and an exogenous (paracrine) one - pro-proliferation. After IR the pro-apoptotic function is predominant: the high apoptotic levels brought about by JNK activity kill the cells shortly after induction thus obliterating the pro-proliferation and pro-migration functions [19]. Previous work has shown that, upon IR or HS, tissues in which cell death is prevented develop overgrowths that require JNK activity [10, 11, 13].

As pointed out above, p53 induction by IR can cause apoptosis independently of JNK, but since both JNK and p53 stimulate each other [20] and both converge at the level of *rpr* and *hid* [15, 17, 19, 26], the overall apoptosis after IR is the result of joint activities of JNK and p53. However, the consequences of the activation of p53 in Apoptosis-Deficient cells (here thereafter referred to as AD cells) have not been investigated.

We have made use of various genetic conditions to study the outcome of subjecting AD tissues to IR. We find that many AD cells acquire high JNK levels and also attain typical features of senescent cells [27]: cellular hypertrophy, cell cycle arrest, oxidative stress, Senescent-Associated (SA) ß-gal activity and Senescent-Associated Secretory Phenotype (SASP). The latter is manifest by the secretion of the signalling molecules Unpaired (Upd), Dpp and Wg, which induce tumorous overgrowth of the surrounding tissue. The acquisition of all these features is completely dependent on JNK activity, for on its absence there is no sign of senescence.

We obtain similar results driving p53 activity in AD tissue, suggesting that the effects caused by IR are mediated by p53 activity. Furthermore, we have analysed the response to IR of AD tissue that is defective in p53 function. The lack of p53 significantly reduces senescence and the magnitude of the overgrowths caused by JNK, indicating that p53 is a major contributor to the senescence induced by JNK.

## RESULTS

### Cells refractory to apoptosis become senescent after irradiation (IR)

For the experiments reported here we have assembled several genetic conditions in which cells cannot execute the apoptosis program: 1) *dronc* mutant cells cannot activate the effector caspases [28]. Larvae mutant for *dronc* develop to late third instar, thus allowing the isolation and manipulation of imaginal discs unable to execute the apoptosis pathway. 2) In discs of genotype *hedgehog-Gal4, tub-Gal80ts* (*hh^Gal80^>) UAS-miRHG* (see full genetic notation in the Methods section) the activity of the three major pro-apoptotic genes *rpr*, *hid* and *grim* is prevented in posterior compartments by the expression of miRNAs that suppress the function of these genes [29]. 3) In discs of genotype *hh^Gal80^>UAS-p35* the presence of the pan caspase inhibitor P35 blocks the activity of the effector caspases in the posterior compartment [30]. Those cells become “undead” and remain alive indefinitely after IR or HS [11, 13].

The activation of the apoptosis program in irradiated AD tissue permits the study of the consequences of unrestrained activity of JNK or p53. We previously reported that IR treatments (4000R) to AD imaginal discs or compartments cause permanent JNK activity, which in turn produces large overgrowths [10]. We have confirmed this observation in discs of genotype *hh^Gal80^>UAS-miRHG*, *TREred* in which apoptosis is inhibited in the posterior compartment and JNK activity monitored by the TREred construct, a reporter of overall JNK activity [31]. After IR, the posterior compartments develop large overgrowths, associated with ectopic and persistent activity of JNK (Extended data Fig. 1a,b and e). The overgrowth of the posterior compartments is strictly dependent on JNK function as expression of *bsk^DN^,* which encodes a dominant negative form of the Jun Kinase Basket suppresses it [32] (Extended data Fig. 1c,d and e).

Given the critical role of JNK in the process, we have paid special attention to the JNK-expressing *TREred* cells. These cells are generally grouped in patches of variable size distributed all over the AD tissue, although they tend to accumulate in the pouch region. On average, the TREred patches fill about 20-25% of the compartment (Extended data Fig. 1b).

The TREred cells also exhibit a number of features, which are essentially like those described for senescent cells (SCs) in *Drosophila* and in vertebrates [27, 33, 34]. 1) Staining with Phalloidin to visualize F-actin shows that TREred cells are bigger than surrounding cells (Fig. 2a,b). 2) TREred cells do not divide, as indicated by the lack of incorporation of cell proliferation markers like EdU (5-ethynyl-2’-deoxyuridine) or the down regulation of the G2/M inducer String (Stg) (Fig. 2c,d). Staining with the Fly-Fucci cell cycle indicator [35] reveals that TREred cells are arrested at the G2 phase of the division cycle (Fig. 2e). 3) TREred cells show high levels of ß-gal activity indicating augmented lysosomal function (Fig. 2f), a typical feature of SCs, referred to as SA-ß-gal activity [27]. 4) They produce Reactive Oxygen Species (ROS) as observed by the activation of the oxidative stress response gene *GstD* (Fig. 2g) [36], another distinctive feature of senescent cells [27]. Moreover, as shown in Fig. 2f,g the gain of ß-gal activity and of ROS production by the TREred cells is strictly dependent on JNK activity.

**Figure 2.**
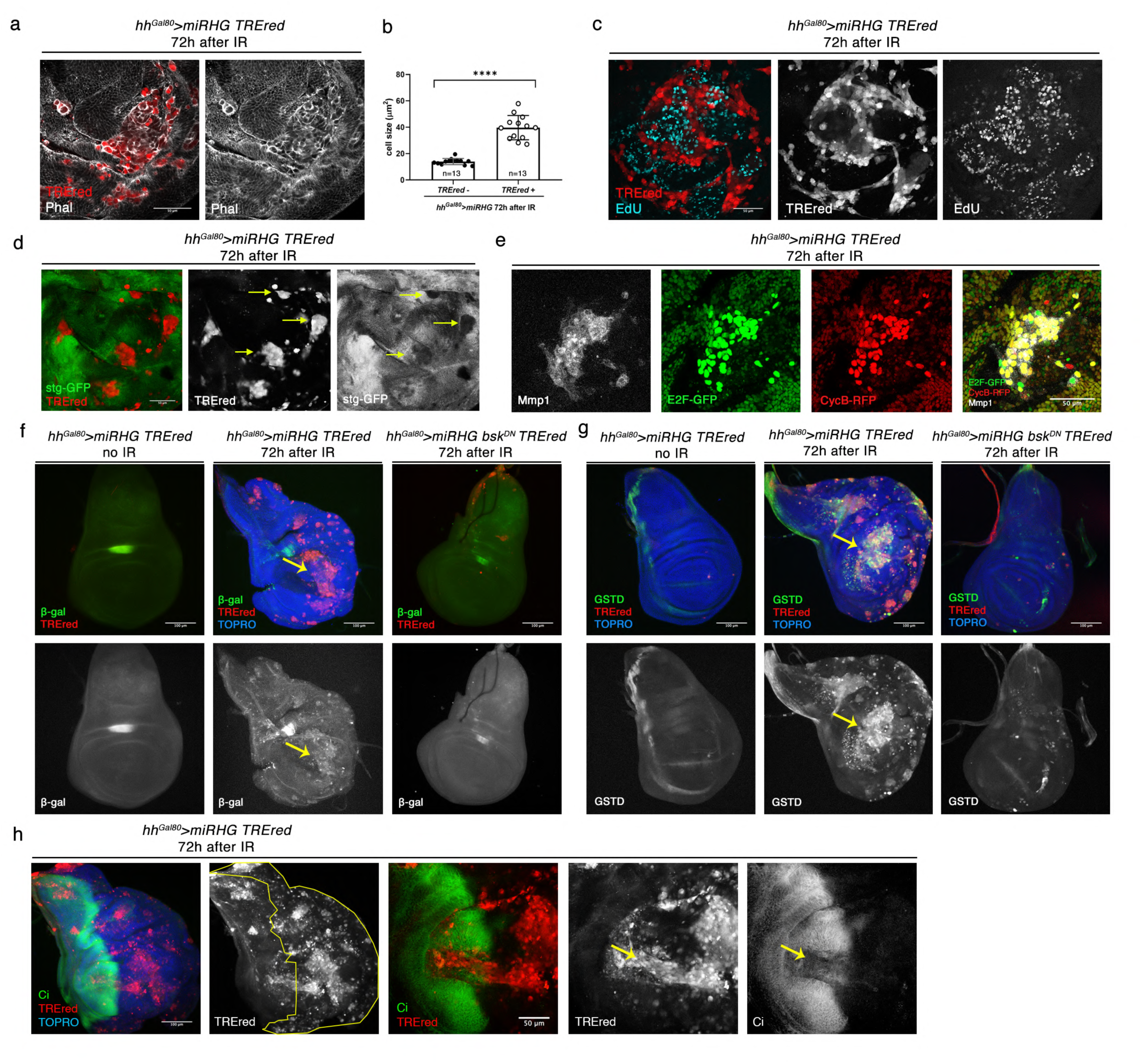
Apoptosis-Deficient cells acquire senescent features after irradiation. The genotypes of the discs and IR treatments are shown on top of the images. a) Fragment of an IR-treated wing disc containing TREred and normal cells. Phalloidin staining (white) reveals the contour of the cells. Those expressing TREred are significantly bigger than the surrounding ones. b) Graph that shows the quantification of the effect on cell size in the genotypes described (n=10 discs and each point is the average cell size of 10 cells). Statistically significant differences based on Student’s t test are indicated: ****=p< 0.0001. c) Double staining for red fluorescence and EdU clearly shows that TREred cells show little or no EdU incorporation in contrast with surrounding cells. The implication is that TREred cells do not divide. d) Double staining for TREred and Stg-GFP shows that Stg is downregulated in the TREred cells, as indicated by the yellow arrows. e) The usage of the Fly-Fucci method allows to determine the phase of the cell cycle in which the JNK-expressing cells, visualised by the expression of the Metalloproteinase1 (Mmp1, a JNK target) are arrested. These cells contain activity of the E2F and CycB degrons, indicating that the JNK-expressing cells are arrested at the G2 period of the cycle. f) Gain of ß-gal activity in TREred cells (yellow arrow), which requires JNK function; there is no gain of ß-gal if JNK is compromised by the expression of a dominant negative form of *basket* (*bsk^DN^*). g) Production of Reactive Oxygen Species (ROS), monitored by the expression of the target gene *GstD*, by the TREred cells. As in the case of ß-gal, it is dependent on JNK activity. h) Invasion of the anterior compartment by TREred cells of posterior origin. The A/P border is delineated by the expression of the *cubitus interruptus (ci)* gene (green), which is expressed only in the anterior compartment. Note that some TREred cells can penetrate a long way into the anterior compartment (yellow arrow). The three images to the left show a high magnification of one penetration into the Ci territory.

We also find a feature not normally associated with senescence, and that is that the TREred cells are able to transgress the antero-posterior (A/P) border and to invade neighbour compartments (Fig. 2h). In the case of the *hh^Gal80^>UAS-miRHG, TREred* experiment we find that in 84% of the discs (n=37) TREred cells, which originate in the posterior compartment, can be found in the anterior one. We have quantified the fraction of the anterior compartment occupied by senescent cells of posterior origin and find it to be around 10%. This indicates a significant contribution to the anterior compartment.

A key feature of senescent cells is the Senescent Associated Secretory Phenotype (SASP); these cells secrete a number of factors, including cytokines, chemokines and metalloproteinases (reviewed in [33]), some of which are associated with inflammation and tumour development in mammalian tissues [37–40]. In our experiments we have analysed whether the TREred cells exhibit secretory phenotype by examining the expression of the genes *upd*, *dpp* and *wg*, which encode the secreted proteins Upd, Dpp and Wg, the ligands of the JAK/STAT, Dpp and Wg pathways respectively. These pathways are known to control growth in imaginal discs [41–43]. As shown in Fig. 3a,b,c, the three genes become ectopically activated in many TREred cells, although not in all. We also found many instances in which there is an increase of the proliferation of cells close to the TREred ones. This effect appeared to be restricted to the central region of the disc (Fig. 3d).

**Figure 3.**
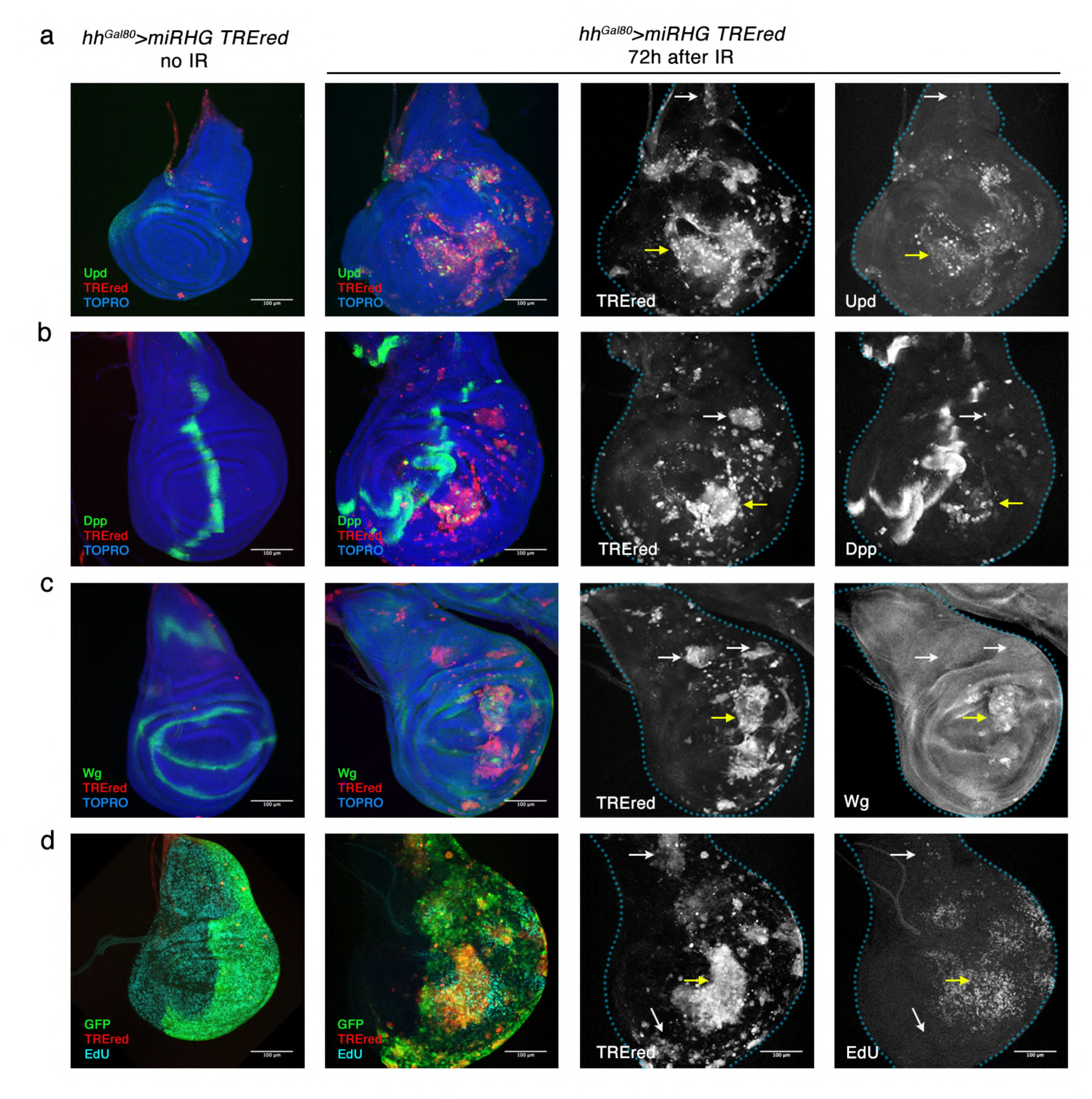
The secretory phenotype of Apoptosis-Deficient cells after irradiation. Comparison of the expression of *upd*, *dpp* and *wg* in control non-irradiated and irradiated discs of genotype *hh^Gal80^>UAS-miRHG, TREred*. a) Ectopic expression of the *upd* gene, monitored by an *upd1-lacZ* marker (green) in TREred cells generated by irradiation. The wildtype expression of *upd1-lacZ* in the control non-irradiated discs is shown. The comparison of TREred and Upd label shown in white in the pictures to the right indicates a high degree of coincidence for both markers (yellow arrow), although not all the TREred cells express *upd* (white arrow). b) Ectopic expression of *dpp*, visualised by a *dpp-lacZ* reporter line (green). The control non-irradiated disc shows the wildtype expression of *dpp*: an anterior stripe along the A/P compartment boundary. As shown in the pictures to the right many TREred cells show ectopic *dpp* expression (yellow arrow), while others do not or at very low levels (white arrow). c) In comparison with the non-irradiated control, many TREred cells express *wg* ectopically in the wing pouch (yellow arrow), although not in the periphery of the disc or the notum region (white arrows). d) Edu incorporation in control and irradiated discs of the genotype *hh^Gal80^>GFP, UAS-miRHG, TREred*. While EdU incorporation (blue) is uniform in the control, the irradiated discs show EdU accumulation in the proximity of the TREred patches located in the central region of the disc (yellow arrow) but not in those in the periphery or in the notum region (white arrows).

### The “undead” cells are senescent

We have also analysed the response to IR and to HS of cells of the posterior compartment of discs of genotype *hh^Gal80^>UAS-p35, UAS-GFP, TREred* in which the presence of baculovirus protein P35 impedes the activity of effector caspases [30] and prevents cell death. This is the same type of experiment in which “undead” cells were described [11, 13]. As illustrated in Extended data. Fig. 2 these undead cells also express senescence biomarkers: they are bigger than surrounding cells, do not divide and produce ROS (Extended data Fig. 2a-d). Moreover, these cells often invade neighbour compartment (Extended data Fig. 2e), a feature we have reported previously [13, 21]. Previous work by us and others showed that these cells secrete the Dpp and Wg ligands [11–13]. We find similar results by generating TREred cells by heat shock (Extended data. Fig. 2f-i). All together, these results demonstrate the senescence nature of undead cells.

### Diversity of SCs cells within the wing disc

In the *hh^Gal80^>UAS-miRHG*, *TREred* experiments above we observed (Fig. 3) that although some senescence biomarkers like size increase or ROS production were present in all TREred cells, there were some TREred cells that did not express the signalling genes *upd*, *wg* and *dpp*. This finding suggested an *in vivo* variability of SCs within the disc, which we have explored in detail. To this effect we irradiated discs of genotype *dronc^-^; TREred,* in which the entire disc is deficient in apoptosis, and examined the expression of senescence biomarkers in TREred patches in the different regions of the disc.

Confirming the preliminary observation, we find two types of TREred patches. One group expresses all the regular senescent biomarkers, e.g. cellular hypertrophy, cell division arrest, SA-ß-gal activity, ROS production (Fig. 4 and Extended data Fig. 3), but do not show expression of the signalling genes *upd*, *dpp* and *wg*. The second group expresses the latter genes in addition to the rest of senescent markers (Fig. 4 and Extended data Fig. 3).

**Figure 4.**
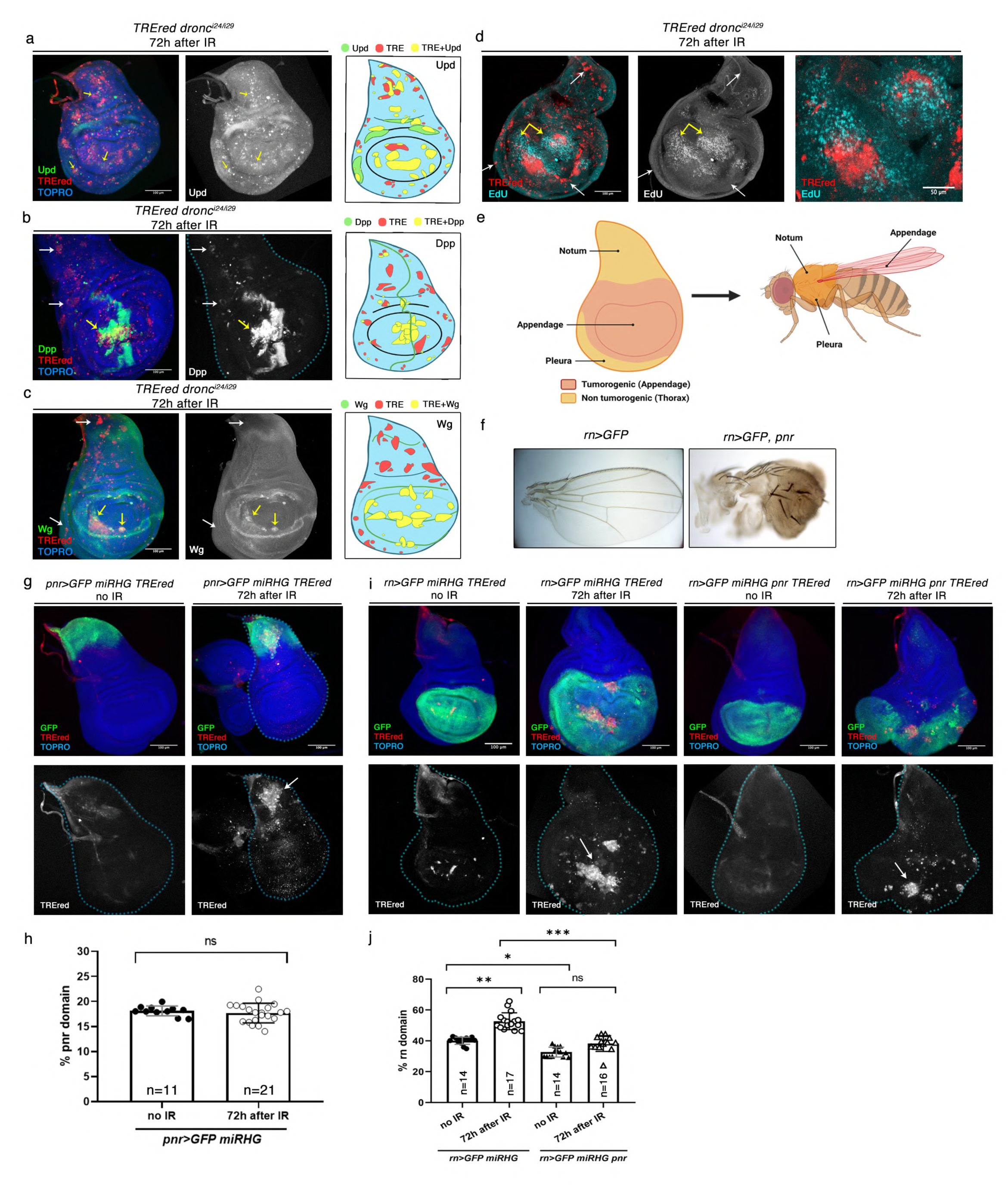
Two types of senescent cells in the wing disc. a) Expression of *upd* (green) in TREred cells in a *dronc* mutant disc after IR, as visualised by the *upd-lacZ* reporter. There is ectopic expression all over the disc (yellow arrows) as illustrated in the scheme to the right which shows the result of a compilation of several discs (n=14). The TREred patches that do not express *upd* appear in red, while those that express *upd* are in yellow. The endogenous expression pattern of *upd* is in green. b) Gain of *dpp* expression (green) by TREred cells in a *dronc^-^* disc after IR. Note that some express *dpp* (yellow arrow) but others do not (white arrows). To the right there is a compilation of several discs (n=13), drawing the TREred patches with *dpp* expression in yellow and the rest in red. It points out the tendency of *dpp*-expressing patches to appear in the wing pouch. The endogenous expression pattern of *dpp* is in green. c) Ectopic expression of *wg* (green) after IR in a *dronc^-^* disc. Note that virtually all the TREred patches expressing *wg* appear in the wing pouch. The result of the compilation of several discs (n=16), clearly illustrates this result: the yellow patches (TREred+Wg) only appear in the wing pouch. The endogenous expression pattern of *wg* is in green. d) EdU incorporation in a *dronc^-^* disc after IR. Note the increase of EdU incorporation (blue) in the vicinity of the TREred patches in the wing pouch (yellow arrows). In contrast, this effect is not observed in the TREred in the periphery of the disc or in the notum region (white arrows). The image to the right shows a high magnification of the pouch region of the disc. e) Summary of the results shown in the (a-d) panels. The wing disc can be subdivided into two regions: the notum and the periphery of the disc, which contain the precursor cells of the thoracic structures (mesothorax and pleura), and the central region, which gives rise to the appendage (wing). As summarized from the results in the (a-d) panels, the tumorigenic region of the disc is restricted to the appendage (red), while the thoracic region is non tumorigenic (orange). f) Transformation of the wing into thorax by driving expression of the *pnr* gene in the wing precursors. In *rn>UAS-GFP, UAS-pnr* flies the wing structures are replaced by thoracic ones, as illustrated (right picture) by the higher density of the trichomes, typical thoracic pigmentation and the appearance of notum-like bristles, which are always absent in the wing appendage (left picture). g) Result of generating senescent cells only in the notum domain. After irradiation discs of genotype *pnr>UAS-GFP, UAS-miRHG, TREred* there is no increase in the size of the Pnr domain (green), although there are numerous TREred cells in the region (white arrow). h) Quantification of the notum domain size in *pnr>GFP, UAS-miRHG, TREred* irradiated and non-irradiated discs represented as % of the Pnr domain respect to the total area of the disc of the disc. n is indicated in the graph. ns, not significant based on Student’s t test. i) This panel displays the results obtained generating senescent cells in the Rn domain (green), which covers the appendage region of the disc. The two images to the left show the overgrowth of the domain after IR, associated with the appearance of TREred cells (white arrow). The two images to the right show the consequences of driving expression of the *pnr* gene in the Rn domain. After IR, in *rn>UAS-GFP, UAS-miRHG, UAS-pnr, TREred* discs there appear numerous TREred cells (white arrow), but there is no overgrowth. j) Quantification of the appendage domain size in *rn>UAS-GFP,UAS-miRHG, TREred* and *rn>UAS-GFP, UAS-miRHG, UAS-pnr, TREred* irradiated and non-irradiated discs represented as % of the Rn domain respect to the total area of the disc. n is indicated in the graph. Statistical analysis by one-way ANOVA when compared the mean of each column with the mean of the control as indicated. *=p<0.05, **=p<0.01, ***=p<0.001 and ns=not significant.

Moreover, the two groups tend to appear in different regions of the disc. As illustrated in Fig. 4a-c, the TREred cells containing *upd*, *wg* and *dpp* expression appear preferentially in the pouch region of the disc, which contains the precursor cells of the appendage (the wing proper). This is more evident in the case of *wg* expression where its activation is restricted to the appendage domain. In contrast, those that do not express *wg* and *dpp* localize mostly to the thoracic domain of the disc, that includes the notum, and the peripheral region of the disc, which gives rise to the pleura, the ventral part of the thorax [44].

Another significant difference between the two groups of SCs is that those in the wing pouch appear associated with over proliferation of neighbour non-SCs, whereas those in the thoracic regions are not (Fig. 4d). We refer to those in the pouch as tumorigenic senescent cells (tSCs), and those in the thorax as non-tumorigenic (ntSCs). We note that the tumorigenic region corresponds very precisely with the appendage part of the disc, whereas the non-tumorigenic region (pleura and notum) corresponds to the thoracic part of the body (Fig. 4e).

We have tested separately the tumorigenic potential of the thoracic and appendage regions of the disc using the *pannier-Gal4 (pnr>)* and the *rotound-Gal4* (*rn>*) lines, which drive expression in the proximal part of the notum and the wing region respectively [45, 46].

In IR-treated discs of genotype *pnr>UAS-miRHG, UAS-GFP, TREred* we observe in the Pnr domain the presence of numerous patches of TREred cells, which show typical senescence features like cellular hypertrophy, lack of cell division, cell motility or SA ß-gal activity (Extended data Fig. 3e-g). However, there is no overgrowth of the Pnr domain, which remains of the same size as the control (Fig. 4g,h). In contrast, *rn>UAS-miRHG, UAS-GFP, TREred* IR-treated discs show a large overgrowth of the Rn domain (Fig. 4i,j)

The different kind of senescent cells in the thorax and the appendage was intriguing; one possibility was that the different local microenvironments of the two regions may influence the type, tSC or ntSC, of the cells, a position-derived effect. Alternatively, it could be determined by the original identity of the cells, thoracic versus appendage, prior to the transformation. To resolve this question, we performed an experiment to change the identity of appendage cells toward thoracic without modifying the position.

It is known that driving expression of the *pnr* gene in wing cells changes their identity from wing to notum [46]. We first confirmed this transformation in adult flies of genotype *rn>UAS-pnr*, which show a clear wing to notum transformation, as indicated by the presence of thoracic bristles and the typical notum pigmentation (Fig. 4f).

Next, we tested whether the change of identity driven by *pnr* in wing cells is reflected in the production of overgrowth, and hence on the type of senescent cell. The finding (Fig. 4i,j) was that unlike the control *rn>UAS-miRHG, UAS-GFP, TREred* discs, which show a clear overgrowth, irradiation of discs of genotype *rn>UAS-miRHG, UAS-GFP, TREred, UAS-pnr* does not cause overgrowth (Fig. 4i). This result suggests that the original identity of the cell is a major factor that determines the type of senescence.

The preceding results demonstrate the existence of at least two classes of SCs in the wing disc, those that secrete the Upd, Dpp and Wg ligands and cause overgrowth in surrounding territory and those that do not. The transformation into tSC or ntSCs appears to be determined by the original identity of the cells.

### Tumorous overgrowths induced by senescent cells; contribution of the JAK/STAT, Wg and Dpp pathways

The presence of SCs in the disc induce the formation of tumorous overgrowths (Extended data. Fig. 1), but detailed examination of division rates indicates that the SCs do not divide, but may stimulate proliferation of non-SCs in their vicinity (Fig. 2C), suggesting that the overgrowths are generated by over proliferating non-SCs.

Moreover, the observation that tSCs generate the Upd, Dpp and Wg growth signals suggests that they induce the activation of the JAK/STAT, Dpp and Wg pathways, which would presumably be responsible for the tissue overgrowth. We have checked this hypothesis in experiments in which the activity of each of those pathways is abolished or reduced in the posterior compartment by specific UAS-RNAi lines. The JAK/STAT pathway is compromised by suppressing the Stat92E transcription factor [41], the Dpp pathway by inactivating the Dpp ligand [47] and Wg by suppressing the function of the receptor Frizzled (Fzr) [48].

Suppression of function of each of the pathways causes a significant decrease in the overgrowth of the posterior compartment, clearly demonstrating their contribution to the tumorigenic process triggered by JNK (Fig. 5a-d).

**Figure 5.**
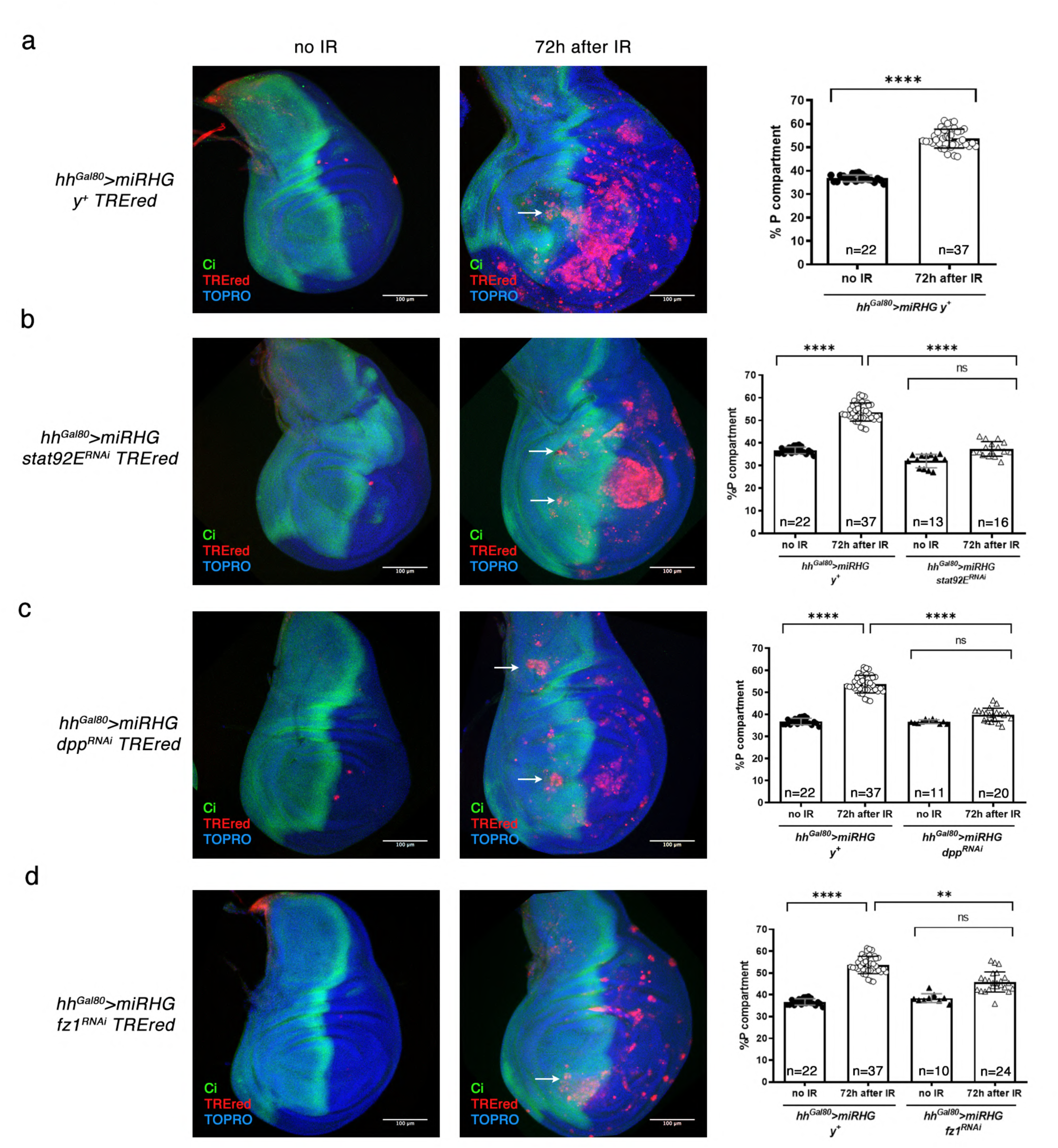
The JAK/STAT, Dpp and Wg pathways are required for the development of overgrowths generated by the senescent cells. a-d) Wing imaginal discs of irradiated and non-irradiated animals of the genotypes described and stained for TOPRO (blue), Ci (green) and TREred (red). White arrow point to TREred cells that can invade the adjacent compartment. a) After IR a disc of the genotype portrayed in the figure develops large overgrowths in the posterior compartment. Discs of this genotype can activate the expression of *upd*, *dpp* and *wg* as seen in Figure 3. The quantification of the magnitude of the overgrowth is shown in the graph to the right. The percentage of the area of the posterior compartment with respect to the total area of the disc is significantly increased with respect to the non-irradiated control. n is indicated in the graph. Statistically significant differences based on Student’s t test are indicated: ****=p<0.0001. b) Effect of eliminating JAK/STAT activity by compromising the function of the *stat92E* gene by RNA interference. After IR the posterior compartment does not overgrow; its size is not significantly different from that of the non-irradiated discs and much smaller than that in a), which contains normal JAK/STAT function. n is indicated in the graph. Statistically significant differences by one-way ANOVA when compared the mean of each column with the mean of the control as indicated. ****=p<0.0001 and ns=not significant. The panels c) and d) show the similar effects of suppressing Dpp (with a *dpp-RNAi* line) or Wg (with a *fzr-RNAi* line that prevents the function of the receptor Frizzled). In neither case there is overgrowth of the posterior compartment, as illustrated by the corresponding graphs. Note that there are fewer TREred cells than in the control in a), but they can still invade the anterior compartment (white arrows). n is indicated in the graph. Statistically significant differences by one-way ANOVA when compared the mean of each column with the mean of the control as indicated. **=p<0.01, ****=p<0.0001 and ns=not significant.

We also find that in the absence of JAK/STAT, Dpp or Wg activity, the amount of TREred cells is significantly reduced (Fig. 5a-d), suggesting a correlation between the number of JNK-expressing cells and the overgrowth of the tissue. It also raises the possibility that the pathways above are required for the maintenance of JNK activity in AD cells. We also note that in spite of the diminution of the number of TREred patches, these can still migrate across compartment borders (Fig. 5). Thus, the lack of signalling does not affect the migratory property of SCs cells.

A principal conclusion from this section is that the JAK/STAT, Dpp and Wg pathways are involved in the development of tumorigenic overgrowths. The specific contributions of each of them to the process and their functional interactions remain to be examined.

### Induction of senescence by p53. Interactions with JNK

Acute activation of p53 after cellular stress has been shown to be implicated in the acquisition of senescence in mammalian cells [49, 50], and also in *Drosophila* [34], but the mechanism by which p53 contributes to senescence in *Drosophila* has not been studied.

It is known that in *Drosophila* p53 is activated after IR, in turn it induces JNK activity and causes an immediate apoptotic response [15, 19]. Thus, the analysis of the non-apoptotic roles of p53 and of its implication in senescence requires the suppression of the apoptotic program in *p53*-expressing cells.

We have tested whether driving expression of *p53* in AD tissue is able to generate senescent cells. To do this, we temporarily expressed *p53* and the *miRHG* construct in the posterior compartment (*hh^Gal80^>UAS-miRHG, UAS-p53*). Posterior cells show strong JNK induction and overgrowth of the compartment (Fig. 6a,b). As in the IR experiments above, TREred positive cells acquire senescence characteristics, including morphology changes as visualized by F-Actin and Topro staining, cell cycle arrest at G2, SA-ß-gal activity, response to ROS and secretory phenotype, as shown by up regulation of *upd*, *dpp* and *wg* (Fig. 6c-i). We also find senescent cells of posterior origin that penetrate in the anterior compartment. These cells contain expression of *dpp* and *wg* (Fig. 6j), what suggests that they affect the growth of the surrounding tissue.

**Figure 6.**
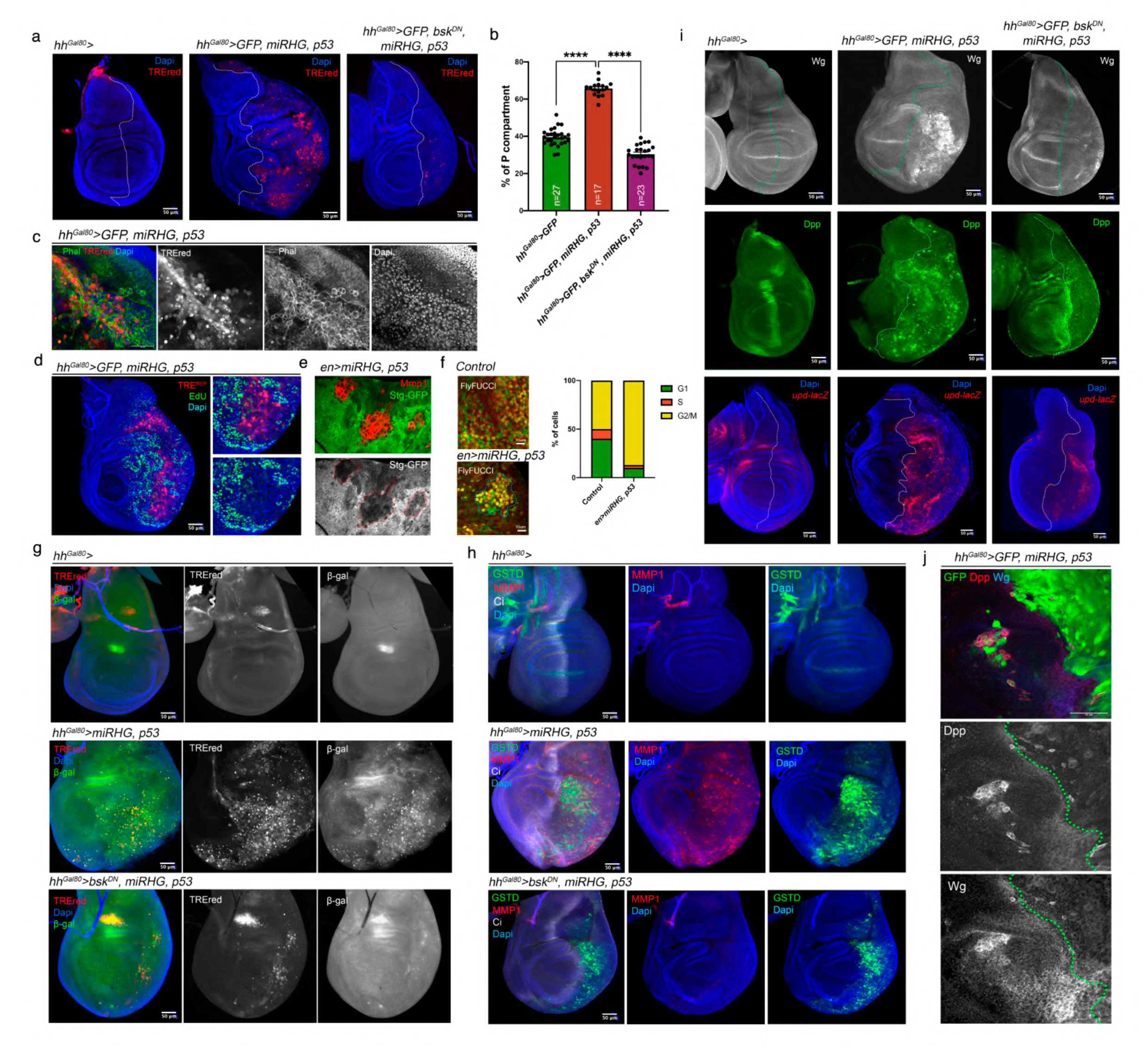
Induction of senescence by p53. The genotypes of the discs are shown on top of the photographs. a) Ectopic expression of *p53* and the *UAS-miRHG* in the posterior compartment for 72 hrs (*hh^Gal80^>*) induces the activity of the TREred and the overgrowth of the compartment. The compartment boundary is indicated as a dotted white line. Dapi is in blue. The overgrowth phenotype as well as the activation of the TREred is supressed by the expression of *bsk^DN^*. b) Quantification of the posterior compartment in the genotypes indicated represented as % of the posterior domain respect to the total area of the disc. n is indicated in the graph. Statistical analysis by one-way ANOVA when compared the mean of each column with the mean of the control as indicated. ****=p<0.0001. c) Fragment of a wing disc containing TREred surrounded by normal cells. Phalloidin staining (green and white) reveals the contour of the cells. Those cells expressing TREred are significantly bigger than the surrounding ones. d) Double staining for TREred and EdU (green) clearly shows that TREred cells show almost no EdU incorporation, in contrast to surrounding cells. The right panels show a magnification of a region of the disc containing TREred cells. e) Double staining for Mmp1 (red) and Stg-GFP (green and white) in an *en>UAS-miRHG*, *UAS- p53,* shows that Stg is downregulated in the red cells, as indicated by the red dotted lines. f) The Fly-Fucci method confirm that Mmp1 cells (staining not shown) are arrested at the G2 phase of the cycle by the presence of the E2F (green) and CycB (red) degrons in control (*en>*) and *p53* expressing discs (*en>UAS-miRHG, UAS-p53*). The quantification of the number of cells that are GFP+, RFP+, and yellow (GFP+RFP+) for each condition is indicated. g) Wing imaginal discs of the genotypes indicated were stained for TREred, Dapi (blue) and ß- gal activity (green). Note that the induction of ß-gal activity by p53 is dependent of the JNK pathway as the expression of *bsk^DN^* strongly abolished. h) Wing imaginal discs of the genotypes indicated were stained for TREred, GSTD (green), Ci (white) and Dapi (blue). i) Ectopic expression of *p53* with the *miRHG* induces the expression of *wg*, *dpp* and *upd* in a JNK dependent manner. Wing imaginal discs of the genotypes indicated were stained for Wg (white), Dpp (green) and *upd-lacZ* (red). The A/P boundary is indicated by a dotted line. j) Magnification of a wing disc of the genotype indicated stained for Wg (red and white), Dpp (blue and white) and GFP (green). A group of *p53, miRHG* expressing cells (green) of posterior origin invade the anterior compartment and activate *dpp* and *wg*. The A/P border is delineated by a dotted line.

Importantly, many of these senescent features brought about by p53, including the ectopic activation of *upd*, *dpp* and *wg* and tissue overgrowth, are suppressed by blocking JNK activity with *bsk^DN^* (Fig. 6a,b,g-i), indicating a functional interaction between p53 and JNK in the induction of senescence.

To analyse the contribution of p53 to the JNK-driven senescence we compared the levels of senescence, measured by the amount of TREred tissue and the size of the overgrown posterior compartments, of irradiated discs of genotypes *hh^Gal80^>UAS-miRHG, TREred, UAS-p53-RNAi*, in which p53 function is abolished and those of genotype *hh^Gal80^>UAS-miRHG, TREred*, which contain normal p53 activity and serves as a control.

The results are shown in Fig. 7. The lack of *p53* function results in a large reduction of the overgrowth of the posterior compartment, as illustrated in Fig. 7a. The size of the posterior compartment with respect to that of the total in irradiated discs decreases from 53% to 42%, while in non-irradiated discs is 38% (Fig. 7b). Furthermore, there is also a large reduction in the amount of TREred tissue in the compartment, from 25 to 8% (Fig. 7c). In non-irradiated controls the amount of TREred tissue is negligible (Fig. 7a). These results clearly demonstrate that much of the cellular senescence caused by JNK is mediated by p53.

**Figure 7.**
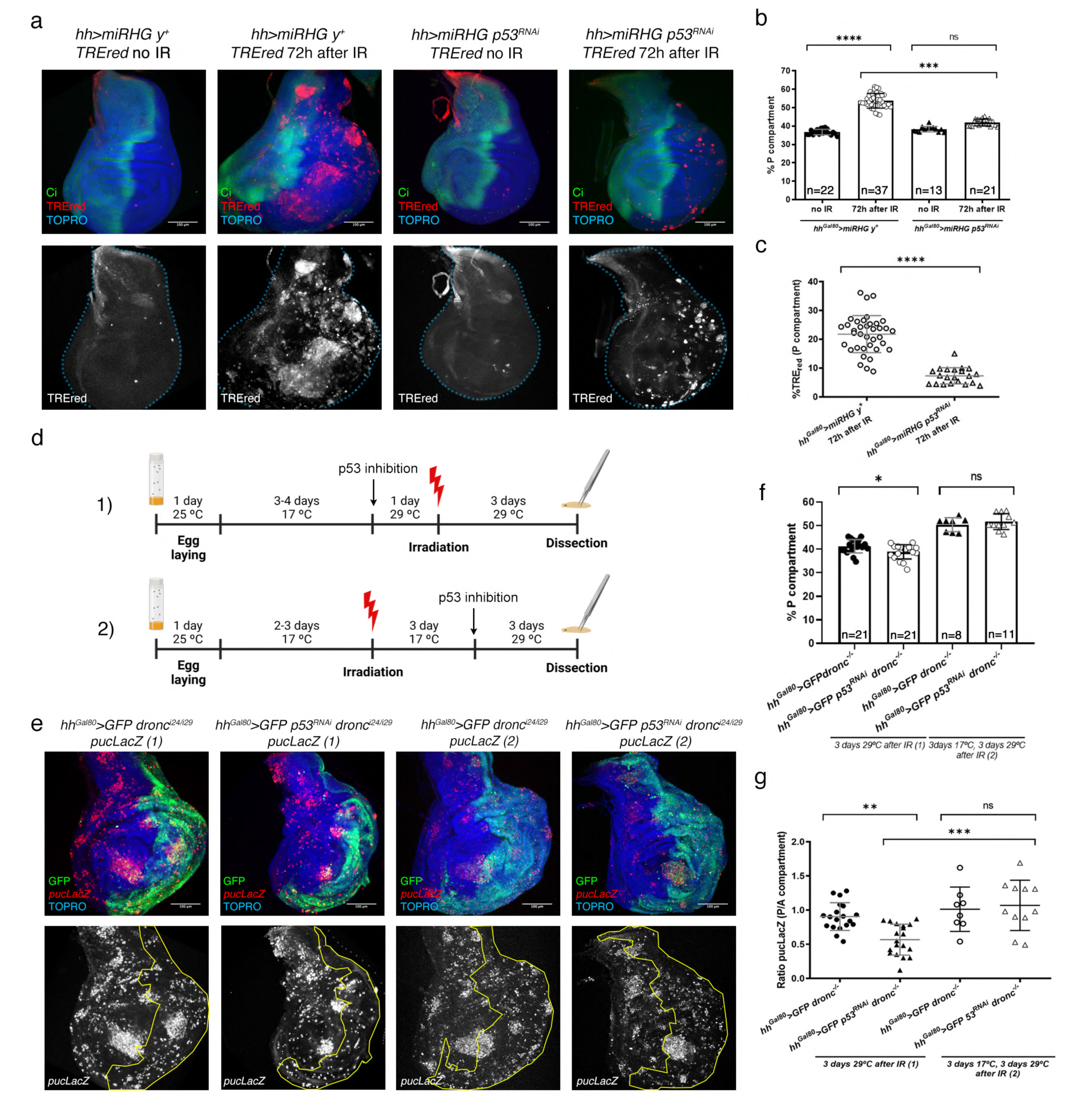
Interactions between p53 and JNK after IR. a) Wing imaginal discs of the genotypes and treatments described above each image stained for TREred (red), TOPRO (blue) and Ci (green) show the effect of the lack of p53 on the levels of senescence, measured by the magnitude of the overgrowth and the amount of TREred tissue. Comparison between the standard *hh^Gal80^>UAS-miRHG, TREred, UAS-y^+^* experiment (control) and the *hh^Gal80^>UAS-miRHG, TREred, UAS-p53^RNAi^* experiment in which p53 function is compromised by RNA interference. In the control experiment we have added an *UAS-yellow* construct (denoted *y^+^* in the figure) so that the number of UAS constructs is the same in the two experiments. This is to ensure there is no titration of the amount of Gal4 product per UAS. After IR, the size of the posterior compartments of the *hh^Gal80^>UAS-miRHG, TREred, UAS- p53^RNAi^* discs is slightly bigger than control non-irradiated discs but the difference is not significant. The number of TREred cells is also much reduced with respect to *h ^Gal80^>UAS- miRHG, TREred* discs, but still higher than in non-irradiated control. b) Quantification of the size of the posterior compartment in the genotypes and treatments indicated represented as % of the posterior domain respect to the total area of the disc. n is indicated in the graph. Statistical analysis by one-way ANOVA when compared the mean of each column with the mean of the control as indicated. ****=p<0.0001 and ns=not significant. c) Quantification of the % of TREred cells in the posterior compartment in the genotypes and treatments indicated. n is indicated in the graph. Statistically significant differences based on Student’s t test are indicated: ****=p<0.0001 and ns=not significant. d) Experiments designed to find the requirement for p53 to generate cellular senescence before and after JNK function is established. The discs are mutant for *dronc* and JNK activity is monitored by the reporter *puc-LacZ*. In experiment (1), larvae of *hh^Gal80^>dronc^-^, UAS-GFP, puc-LacZ* (control) and *hh^Gal80^>dronc^-^, UAS-GFP, puc-LacZ, UAS-p53^RNAi^* genotype are grown for 3-4 days at 17°C before temperature shift to 29°C to suppress p53 function in the posterior compartment of the experimental discs. After one day at 29°C the larvae are irradiated and subsequently dissected 3 days later. In experiment (2), larvae of the same genotypes were irradiated to induce JNK activity and the temperature shift was administered 3 days after the irradiation to allow JNK activation before p53 downregulation. e) The four images to the left show the results of experiment (1) and to the right those of experiment (2). The suppression of p53 in the posterior compartment before irradiation in experiment (1) reduces the number of *puc-LacZ* cells and prevents the overgrowth of the compartment, which is significantly smaller than the control with normal p53 function, as shown in the graph in f. In contrast, the results of experiment (2) indicate that the suppression of p53 after JNK is active has no effect on the induction of *puc-LacZ* cells and on growth of the posterior compartment. The posterior compartment is visualized by GFP and in the images below as a yellow contour. f) Quantification of the size of the posterior compartment in the genotypes and treatments indicated represented as % of the posterior domain respect to the total area of the disc. n is indicated in the graph. Statistically significant differences based on Student’s t test are indicated: *=p<0.05 and ns=not significant. g) Quantification of the ratio of *puc-lacZ* staining between the posterior and anterior compartment of the genotypes and treatments described in (d) and (e). Statistically significant differences by one-way ANOVA when compared the mean of each column with the mean of the control as indicated. **=p<0.01, ***=p<0.001 and ns=not significant.

However, is it possible that p53 could be also involved in the mechanism maintaining the permanent JNK activity observed in AD cells after its initial activation. To check on this possibility we performed experiments to suppress p53 after JNK activity is established (Fig. 7d). In irradiated discs of genotype *hh^Gal80^>UAS-p53-RNAi, UAS-GFP, dronc^-^* we inactivated p53 in the posterior compartment 72 hrs after IR, well after the initial JNK activation (Fig. 7d,e). Then, we examined the effect of the suppression of p53 by comparing the size of the posterior compartments with or without p53 function (Fig. 7e,f). The result is that the suppression of p53 activity after JNK function has been established has no detectable effect on the size of the posterior compartment (Fig. 7f), nor on JNK activity, as measured by the expression of the JNK target gene *puckered* (*puc*) [51] (Fig. 7g). These findings indicate that p53 only contributes to senescence by enhancing the initial activation of JNK.

## DISCUSSION

### Acquisition of senescence by *Drosophila* Apoptosis-Deficient (AD) cells after IR or p53 induction

We have performed several experiments subjecting *Drosophila* AD cells, which cannot execute the apoptosis program, to treatments that would normally result in massive cell death: a high IR dose (4000R) or induction of the *p53* gene, which triggers apoptosis. In those experiments, a fraction of the AD cells acquires high and persistent levels of JNK activity, visualized by the TREred reporter. In addition, they exhibit the typical attributes of senescence, such as cellular hypertrophy, cell cycle arrest, SA ß-gal expression, ROS production and the secretory phenotype (SASP) (see reviews in [33, 37, 52]. Our conclusion is that cells in which the apoptosis program is inhibited are prone to become senescent.

In addition to the biomarkers described above, we have also observed that SCs possess migratory behaviour, that can be visualized by invasion of neighbour compartments, which may extend to long distance from the A/P border. This behaviour is observed in irradiated *hh^Gal80^>UAS-miRHG, TREred* (Fig. 2h) and *hh^Gal80^>UAS-GFP, UAS-p35, TREred* discs (Extended data Fig. 2e), as well as after overexpressing *p53* in *hh^Gal80^>UAS-GFP, UAS-miRHG, UAS-p53* (Fig. 6j). This invasiveness property has also been reported for apoptosis-deficient aneuploid cells [24], which become senescent [53]. The ability to migrate is dependent on the activation of the JNK transcription program and can be reversed by suppressing JNK [24]. Although in the latter report the authors associate invasiveness to aneuploidy, our experiments suggest that invasiveness does not depend specifically on aneuploidy but it is a general property of SCs, which express one of the non-apoptotic roles of JNK activity, the stimulus of cell migration. This property has also been reported in mammalian cells expressing JNK [23] [22] and during the normal development of *Drosophila* [25, 54]. In our experiments this property is common to tumorigenic and non-tumorigenic SCs. The ability to invade neighbour tissues may also be a general biomarker of senescence and one that might have implications in tumorigenesis. Regarding tumorigenesis, it is of interest that the migratory property of senescent cells has already been shown to be associated with metastasis in thyroid cancer in humans [55].

Our results also bear on one feature, resistance to apoptosis, generally considered diagnostic of SCs (see [27]). Our experiments suggest that, at least in *Drosophila*, resistance to apoptosis may not be a consequence of senescence but a prerequisite to become senescent. In tissues open to apoptosis, IR treatments or *p53* expression do not induce cells to become senescent; these are rapidly eliminated by the pro-apoptotic function of JNK and p53. In contrast, in tissues deficient in apoptosis those treatments result in permanent expression of JNK, which drives cells into acquiring the typical attributes of senescence. We propose that it is not that SCs avoid apoptosis but that cells refractory to apoptosis are prone to become senescent.

There have been several previous reports describing senescence in *Drosophila* imaginal cells. Nakamura et al 2014, showed that mitochondrial dysfunction associated with Ras activation causes senescence in eye imaginal cells, with features like those described above [34]. The induction of senescence appears to be mediated by production of ROS in the affected cells. This results in p53 and JNK activity, which appear to be responsible for the senescent phenotype.

We note, however, that in our experiments SCs are arrested in G2, whereas in Nakamura et al 2014 the up regulation of the *p21* ortholog *dacapo* indicates cell cycle arrest at the G1 stage [34]. Whether this difference is significant regarding senescence is not clear, but it indicates that *Drosophila* imaginal SCs may use different mechanisms to arrest cell divisions.

Another example of senescence in *Drosophila* has been provided by Joy et al 2021 [53]. These authors induce chromosomal instability by inactivating the spindle assembly checkpoint gene *rod*, what causes aneuploidy. To circumvent the poor viability of aneuploid cells, they are generated in an apoptosis-deficient background, using the miRHG construct, which inactivates the *rpr*, *hid* and *grim* pro-apoptotic genes [29]. Under these genetic conditions aneuploid cells survive and display typical senescence biomarkers. Importantly, the acquisition of senescence is dependent on JNK activity.

### p53 in senescence

The implication of p53 in the acquisition of senescence in mammalian cells is well established (reviewed in [49, 50]). Cellular stress like IR leads to p53 activation and the transcriptional induction of many target genes, some of which, *p21* for example, are effectors of senescence biomarkers like cell cycle arrest.

In the imaginal discs of *Drosophila,* p53 activation by IR induces the pro-apoptotic genes, which results in cell death in a p53-dependent manner [17] [18] (reviewed in [15]). However, here we show that in abscence of apoptosis, p53 function drives cells into senescence (Fig. 6). The involvement of p53 has also been demonstrated in senescence induction by Ras overexpression in eye imaginal discs [34].

In comparing the roles of JNK and p53 as effectors of senescence, the key observation is that there is no senescence in the absence of JNK (Fig. 2 and 6), while in the absence of p53 senescence is reduced but not abolished (Fig. 7a). Thus, the pro-senescence role of p53 requires JNK function, suggesting that p53 exerts a subsidiary role, likely enhancing JNK activity. Furthermore, this enhancing function of p53 appears to act only during the initial activation of JNK, for once it is established the loss of p53 activity does not affect the levels of senescence (Fig. 7d-g).

### A model of senescence in *Drosophila*

Based on previous work and on our own results we propose that there are two critical prerequisites for *Drosophila* imaginal cells to become senescent. 1) inhibition of apoptosis, 2) a trigger of JNK activity (IR, p53 and likely other stressors). Under these conditions the pro-apoptotic function of JNK is suppressed, the AD cells remain alive and manifest the non-apoptotic roles of JNK. The initial triggering of JNK is resolved into permanent activity due to a maintenance loop involving *moladietz* and ROS [10, 56].

This proposal provides a frame to integrate the published results about senescence in *Drosophila*. The aneuploid apoptosis-deficient cells reported by Joy et al 2021 produce ROS and subsequent activation of JNK, which becomes permanent; in our view the key issue is that the process producing aneuploidy also activates the JNK pathway, which is responsible for the senescence.

In the experiments by Nakamura et al 2014 we believe that a primary reason for the transformation is that the elevated levels of Ras activity make the cells refractory to apoptosis [10, 57, 58]. Then the stress induced by the mitochondrial dysfunction activates JNK, which becomes permanent in the absence of apoptosis. We have previously shown [10] that a simple IR stress administered to cells expressing *Ras^V12^* causes ectopic long-term JNK activity associated with *wg* induction and massive overgrowths; a senescence scenario.

The migratory property of SCs described in this report and in previous work [12, 13, 24] is also consistent with the proposal, since one the cellular functions associated with JNK is the induction of cell motility [23] [22, 25].

### Induction of tumorigenesis: the tSC and ntSC senescent cells. Role of identity

The secretory phenotype is of special interest because it is responsible for paracrine effects of senescence like tumorigenesis. We have identified three secreted signals, Upd, Wg and Dpp, although surely there will be others. Interestingly, unlike other senescence biomarkers, the Upd, Wg and Dpp ligands are not expressed in all the SCs cells, but only in a fraction of about 50%. Moreover, only the SCs expressing those ligands induce non-autonomous overgrowths (Fig. 3d, Fig. 4d).

This reveals a genetic diversity within the wing disc (Fig. 4a-e): after IR there are two classes of SCs: tSCs, which produce and secrete proliferative signals and generate overgrowths and ntSCs, which do not. Although genetic variability has been described for senescent cells in culture [59], to our knowledge this is the first case reporting this kind of variability of SCs in *in vivo* tissues.

Of special interest is that the tSC and ntSCs localize to different regions of the disc (Fig. 4a-c,e). The former appears preferentially in the appendage region while the ntSCs appear in the trunk region (notum and the pleura). The implication is that, regarding senescence, the wing disc is subdivided into a tumorigenic and a non-tumorigenic region. Strong support for this view is our finding that generating SCs exclusively in the thorax domain does not cause tumorigenic overgrowth (Fig. 4g,h). In contrast, induced SCs in the wing pouch produce large overgrowths (Fig. 4i,j).

An important question is about the mechanism behind this position-related diversity. Our experiments suggest that a critical factor is the original identity of the cells prior to the transformation towards senescence. Cells with appendage (wing pouch) identity can generate overgrowths after IR (Fig. 4i), but those with thoracic (notum and pleura) identity do not (Fig. 4g). This idea is strongly reinforced by the experiment in which cells of the wing pouch are transformed into notum cells and consequently lose to ability to generate overgrowth (Fig. 4f-j).

Since JNK is the key factor inducing senescence, the implication is that it functions differently in cells of thoracic and appendage identity. In fact, there is previous evidence of the differential activity of JNK in the two regions: driving a constitutive active form of *hemipterous* (*hep^CA^*), the effector of JNK [32] in the wing pouch induces overgrowth, while it does not in the notum region [21]. A finding likely to be related with the inability of thoracic structures to regenerate after massive damage [21]. The distinct behaviour of SCs in the thorax and the appendage may well be a reflection of the very different patterning and genetic mechanisms operating in the body trunk and the appendages [60].

The genetic diversity of SCs we find in *Drosophila* impinge on a classical paradox of the role of senescence: its anti-tumour and pro-tumour properties, and associate them with the identity of the cells prior to the acquisition of the senescence. The anti-tumour role is expressed by the senescent cells of thoracic identity (ntSCs); they stop dividing and do not activate the Wg and Dpp pathways. In contrast, the senescent cells of appendage identity (tSCs, located in the wing) produce the Upd, Dpp and Wg ligands that induce activity of these pathways, known to stimulate growth and proliferation of neighbour cells. The persistent activity of those pathways causes excess of growth: a pro-tumour role.

## Methods

### *Drosophila* strains

The different *Drosophila* strains were maintained on standard medium at 25 °C (except for the temperature shift experiments, see below). The strains used in this study were the following:

Gal4 lines: *hh-Gal4* ([61]), *pnr-Gal4* ([62]), *rn-Gal4* ([45]), *tub-Gal80^ts^* ([63]) and *en-gal4* (Flybase ID: FBti0003572).

UAS lines: *UAS-miRHG* ([29]), *UAS-y^+^* ([62]), *UAS-bsk^DN^* (Bloomington Drosophila Stock Center, BDSC#6409)*, UAS-GFP* (BDSC), *UAS-Stat92E^RNAi^* (BDSC#33637), *UAS-p53^RNAi^* (BDSC#29351), *UAS-pnr* ([64]), *UAS*-*p53-A* ([65]), *UAS*-*dpp^RNAi^* ([66]), *UAS-fz1^RNAi^* (VDRC#105493).

Mutants: *dronc^i29^*, *dronc^i24^* (a gift from A. Bergmann, MD Anderson Center, Houston, TX, USA)

Reporter lines: *TREred* ([31]), *stg-GFP* (BDSC#50879), u*bi-GFP-E2f1^1-230^, ubi-mRFP1-NLS-CycB^1-266^* (Fly-Fucci, BSDC#55099), *GstD-LacZ* ([36]), *upd1-LacZ* (a gift from T. Igaki), *dpp-LacZ* (p10638, BDSC#12379), *puc-LacZ (puc^E69^)* ([51]).

### Imaginal discs staining

Third instar larvae were dissected in PBS and fixed with 4% paraformaldehyde, 0.1% deoxycholate (DOC) and 0.3% Triton X-100 in PBS for 27 min at room temperature. They were blocked in PBS, 1% BSA, and 0.3% Triton, incubated with the primary antibody over night at 4 °C, washed in PBS, 0.3% Triton and incubated with the corresponding fluorescent secondary antibodies for at least 2 h at room temperature in the dark. They were then washed and mounted in Vectashield mounting medium (Vector Laboratories).

The following primary antibodies were used: rat anti-Ci (DSHB 2A1) 1:50; mouse anti- Mmp1 (DSHB, a combination, 1:1:1, of 3B8D12, 3A6B4 and 5H7B11) 1:50; mouse anti-β-galactosidase (DSHB 40-1a) 1:50; rabbit anti-β -galactosidase (ICN Biomedicals) 1:2000; mouse anti-Wingless (DSHB 4D4) 1:50; rabbit anti-Dpp ([67]) 1:200; rabbit and mouse anti-PH3 (MERCK and Cell Signal Technology) 1:500, rat anti-RFP (Chromotek, 5F8) 1:2000.

Fluorescently labelled secondary antibodies (Molecular Probes Alexa-488, Alexa-555, Alexa-647, ThermoFisher Scientific) were used in a 1:200 dilution. Phalloidin TRITC (Sigma Aldrich) and Phalloidin-Alexa-647 (ThermoFisher Scientific) were used in a 1:200 dilution to label the actin cytoskeleton. TO-PRO3 (Invitrogen) and DAPI (MERCK) was used in a 1:500 dilution to label the nuclei.

### EdU incorporation

Dissected wing discs were cultured in 1 mL of EdU labelling solution for 20 min at room temperature and subsequently fixed in 4% paraformaldehyde for 30 min at room temperature. Rat anti-RFP (Invitrogen) 1:200 antibody was used overnight at 4 °C before EdU detection to protect RFP fluorescence. EdU detection was performed according to the manufacturer instructions (Click-iT EdU Alexa Fluor 647 Imaging Kit, ThermoFisher Scientific), and wing discs were incubated during 30-40 min at room temperature in the dark.

### β-galactosidase activity assay

To assess the levels of β-galactosidase activity in senescent cells a CellEvent^TM^ Senescence Green Detection Kit (Invitrogen) was used. Larvae were fixed in 4% paraformaldehyde for 20 min and, after washing with PBS, 1% BSA, incubated with the Green Probe (1:1000 dilution) for 2 h at 37ªC in the dark. Larvae were washed with PBS and then wing imaginal discs were mounted in Vectashield. Images were taken shortly after the protocol to avoid loss of fluorescence.

### IR treatments

Larvae were irradiated in an X-ray machine Phillips MG102 at the standard dose of 4000Rads (R) at different times after the egg laying and dissected at the indicated times depending on the experiment (see “Temperature shift experiments”).

### Temperature shift experiments

Generation of senescent cells in posterior compartments: larvae of the genotype *hh- Gal4, tub-Gal80^ts^* (*hh^Gal80^>*); *TREred, UAS-miRHG* were raised at 17°C for 3-4 days and then transferred to 29°C 1 day before irradiation to activate the corresponding transgene. After irradiation, larvae were kept at 29ªC for 3-4 days to allow tumor growth before being dissected. This standard protocol was used to analyse all senescent markers as well as the contribution of JNK, p53, JAK/STAT, Dpp and Wg pathways to senescence and size of the compartment. This protocol was also used for larvae of the genotypes *hh-Gal4, tub-Gal80^ts^; TREred; UAS-p35*.

Induction of *p53-A* and *miRHG* in the posterior compartment: larvae of the genotype *hh-Gal4, tub-Gal80^ts^*; *TREred, UAS-miRHG; UAS-p53-A* were raised at 17°C for 5-7 days and then transferred to 29°C for 3 days before dissection to activate the corresponding transgene. This standard protocol was used to analyse all senescent markers as well as to study the contribution of JNK. For the Fly-Fucci and Stg-GFP experiments, the *en-Gal4* line was used and the larvae were kept at 25°C before dissection.

Generation of senescent cells in *dronc* mutant discs: larvae of the genotype *TREred; dronc^i24^/dronc^i29^* were raised at 25°C and changed to 29°C 1 day before irradiation. After IR treatment, larvae were kept at 25°C for 3-4 days before dissection.

Generation of senescent cells in the Pnr and Rn domains: larvae of the genotype 1) *TREred, UAS-miRHG/UAS-GFP; pnr-Gal4*, 2) *TREred, UAS-miRHG/UAS-GFP; rn-Gal4* were raised at 25°C and changed to 29°C 1 day before irradiation. After IR treatment, larvae were kept at 25°C for 3-4 days before dissection.

Generation of *pnr*-expressing cells in the Rn domain: Larvae of genotype *TREred, UAS-miRHG/UAS-GFP; rn-Gal4/UAS-pnr* were raised at 25°C and changed to 29°C 1 day before irradiation. After IR treatment, larvae were kept at 25°C for 3-4 days before dissection.

Experiment to analyse the contribution of p53 on the JNK-induced senescence: Larvae of the genotypes 1) *UAS-GFP, tub-Gal80^ts^; hh-Gal4 dronc^i29^/dronc^i24^, puc-LacZ* and 2) *UAS-GFP, tub-Gal80^ts^/UAS-p53^RNAi^; hh-Gal4 dronc^i29^/dronc^i24^, puc-LacZ* were subjected to two different experimental treatments. The first one was the same as described above for the generation of senescent cells in posterior compartments; in the second one, larvae were raised at 17°C for 2-3 days, irradiated and kept at 17°C for 3 days to allow normal JNK activation. After this, larvae were changed to 29°C for 3 days to allow expression of the construct *UAS-p53-RNAi* and then dissected. An illustrative scheme of the two experimental treatments is presented in Fig.7D.

### Image acquisition, quantifications and statistical analysis

Stack images were captured with a Leica (Solms, Germany) LSM510, LSM710, DB550 B vertical confocal microscope and a Nikon A1R. Multiple focal planes were obtained for each imaginal disc. Quantifications and image processing were performed using the Fiji/ImageJ (https://fji.sc) and Adobe Photoshop software. Schemes in Fig.1, Fig.4 and Fig.7 were made with BioRender (https://www.biorender.com/). For quantification of the percentage of the posterior compartment/domain, a Z-maximal intensity projection was made for each image. Then, the area of the compartment/domain (labelled with GFP or the absence of Ci staining in each case) was measured by using the “Area” tool and normalized dividing between the total disc area (labelled by TOPRO-3 or DAPI staining). For JNK activation, quantification *(%TREred*), *TREred* positive area was measured using the “Threshold” tool and then normalized dividing between the area of the compartment. *pucLacZ* ratio (P/A) is calculated as the *%pucLacZ* in the posterior compartment divided between the *%pucLacZ* in the anterior compartment. Cell size was measured as the mean of the area of 10 *TREred* positive cells (labelled by phalloidin staining) and the mean of the area of 10 *TREred* negative cells per imaginal disc.

For the Fly-FuccI cell quantification, third instar larvae of the *en-Gal4; UAS-p53-A, UAS-miRHG; UAS-GFP-E2F1^1-230^, UAS-mRFP1-NLS-CycB^1-266^* genotype were raised at 25°C and stained for Mmp1 and the Fly-Fucci markers. Red, green, or yellow cells were manually quantified. For each experiment, at least five wing imaginal discs were used to count at least 50 Mmp1 positive group of cells.

Statistical analysis was performed using the GraphPad Prism software (https://www.graphpad.com). To compare between two groups, a non-parametric Student’s *t*-test test was used. To compare between more than two groups, a non-parametric, one-way ANOVA Dunnett’s test was used. Sample size was indicated in each Figure legend.

## Acknowledgements

We thank Brian Calvi, Marco Milán, the Bloomington Stock Center, the Vienna Drosophila Resource Center, and the Developmental Studies Hybridoma Bank for fly stocks and reagents. Thanks also to the Confocal Microscopy Service at CBMSO for their help. We especially thank Ernesto Sanchez-Herrero, Antonio Baonza, Mireya Ruiz-Losada and the members of the Morata lab for fruitful discussions throughout the period of this work.

This study was supported by grants from: FEDER/Ministerio de Ciencia e Innovación-Agencia Estatal de Investigación-Consejo Superior de Investigaciones Científicas [No. PGC2018-095151-B-I00, PID2021-125377NB-100, PIE Intramural 202020E255 to GM and No. PGC2018-095144-B-I00, PID2021-127114NB-100 to CE).

---

**Extended data Figure1.**
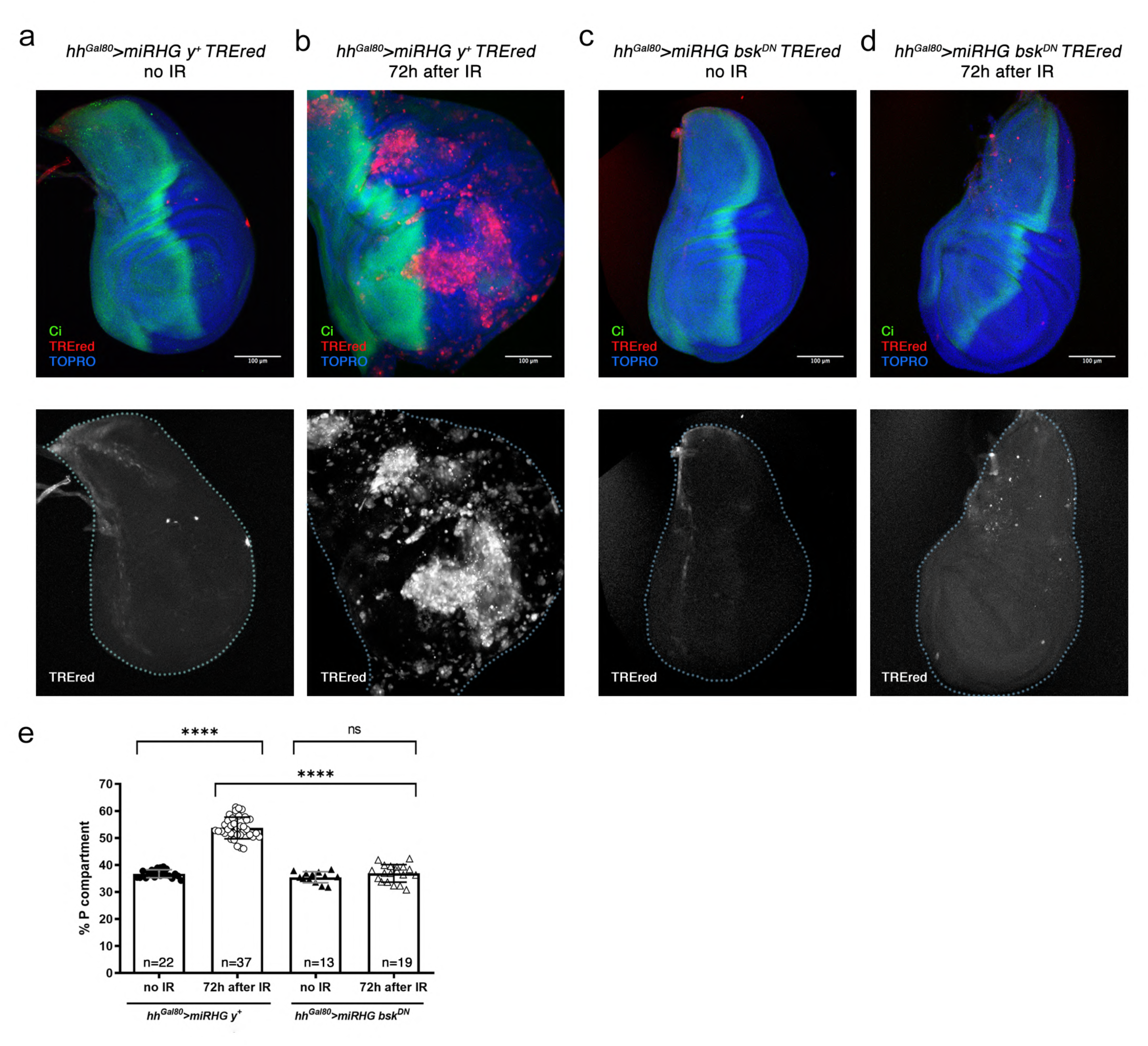
Effect of irradiation (4000R) on apoptosis-deficient posterior compartment and role of JNK. Wing imaginal discs of the genotypes and treatments described above each image stained for TREred (red), TOPRO (blue) and Ci (green). a) Non-irradiated disc of *hh^Gal80^>UAS-miRHG, TREred* phenotype shown normal size and virtually no cells expressing TREred. b) Disc of the same genotype as in (a) 72h after IR. Note the significant increase of size of the posterior compartment, quantified in (e), and the accumulation of cells expressing the TREred construct. c-d) The panels illustrate the response to IR of discs in which the posterior compartments cells contain a dominant negative form of the kinase Basket (*bsk^DN^*), which makes the JNK pathway ineffective. Note in (d) that the IR has no effect on discs of this genotype. e) Quantification of the size of the posterior compartment in the genotypes and treatments indicated, represented as % of the posterior domain respect to the total area of the disc. n is indicated in the graph. Statistical analysis by one-way ANOVA when compared the mean of each column with the mean of the control as indicated. ****=p<0.0001 and ns=not significant.

**Extended data Figure 2.**
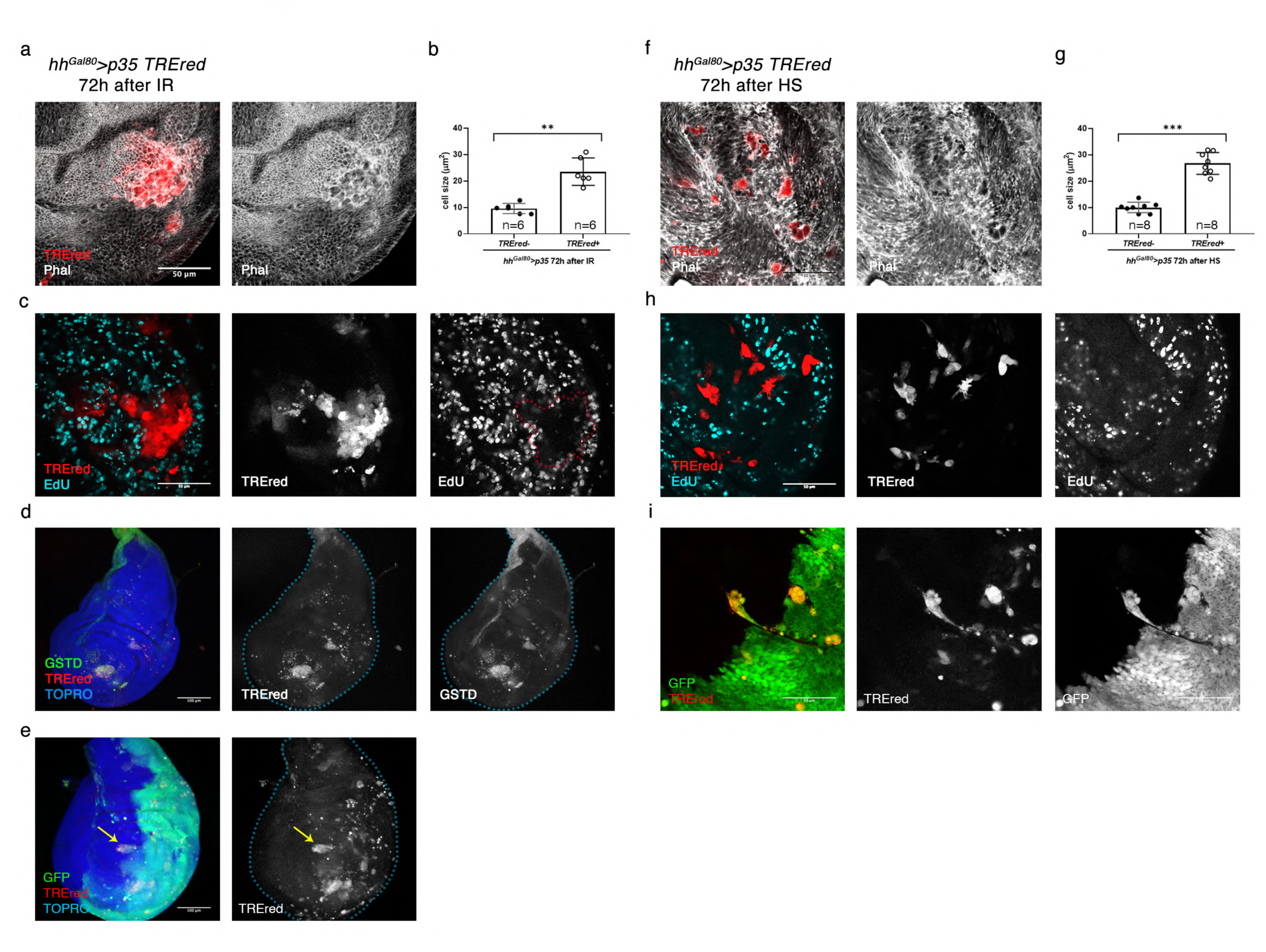
The “undead” cells are senescent. a) Part of a wing disc of genotype *hh^Gal80^> p35, TREred* after 4000R IR. Phalloidin (white) staining reveals that the size of TREred cells is bigger than that of surrounding cells. b) Graph that shows the quantification of the effect on cell size in the genotypes described (n=6 discs and each point is the average cell size of 10 cells). Statistically significant differences based on Student’s t test are indicated: **<p=0.01. c) EdU incorporation (blue) shows that TREred cells do not divide, in contrast to surrounding cells. d) Wing disc of genotype *hh^Gal80^>UAS-p35, TREred* after 4000R IR. TREred cells also show expression of the *GstD-lacZ* (green), a reporter of ROS production. e) Invasion of the anterior compartment by TREred cells of posterior origin (yellow arrow). f) TREred cells generated after 3h heat shock (HS) at 37°Cof the genotype *hh^Gal80^>UAS-p35, TREred* become bigger that surrounding cells, as revealed by Phalloidin staining (white). g) Graph that shows the quantification of the effect on cell size in the genotypes described (n=8 discs and each point is the average cell size of 10 cells). Statistically significant differences based on Student’s t test are indicated: ***=p<0.001. h) EdU incorporation (blue) shows that TREred cells do not divide, in contrast to surrounding cells in *hh^Gal80^>UAS-p35, TREred* after 72h of HS. i) Invasion of the anterior compartment by TREred cells of posterior origin in *hh^Gal80^>UAS-p35, UAS-GFP, TREred* wing discs after 72h of HS.

**Extended data Figure 3.**
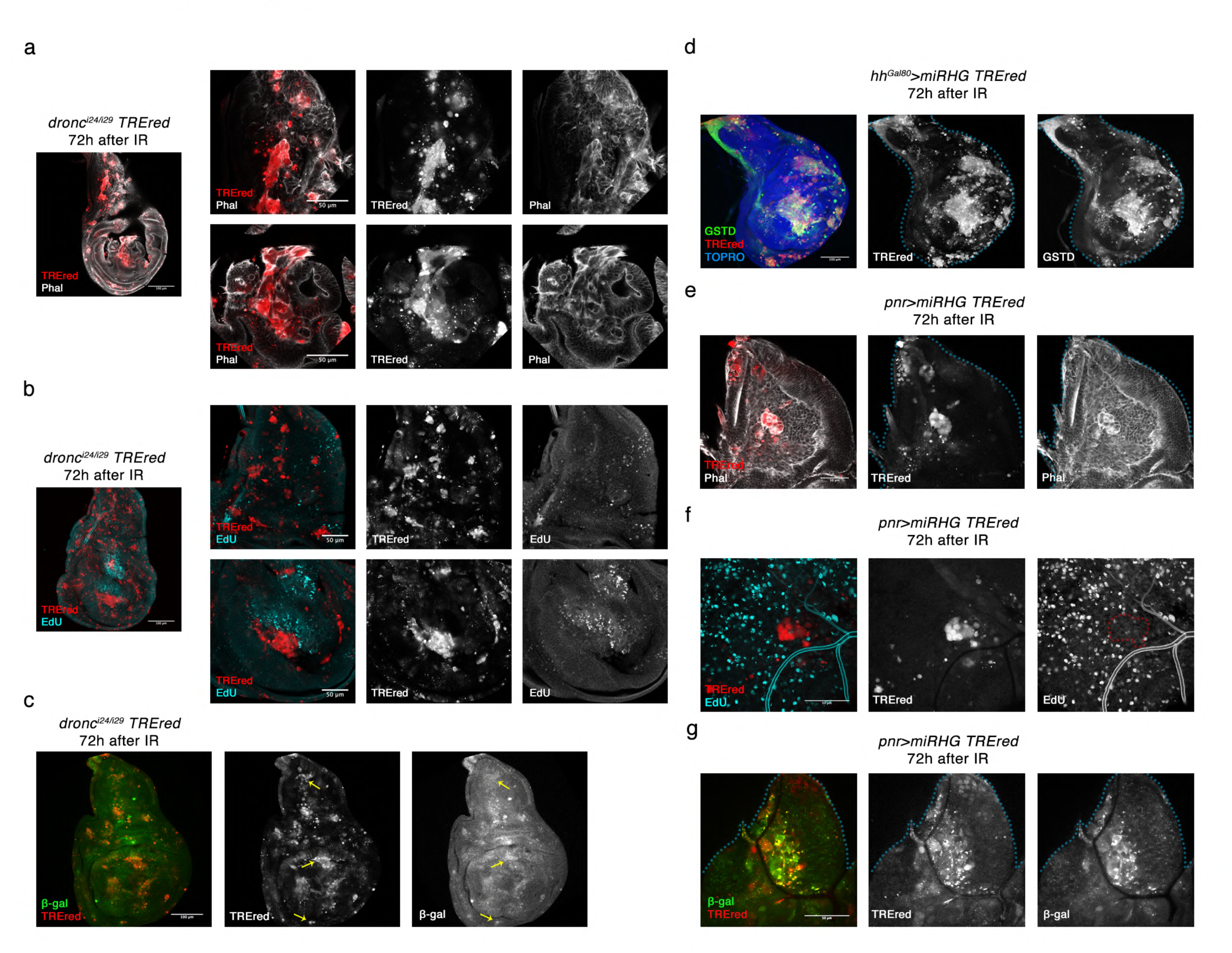
Senescent features in *dronc^-^* discs and in the notum region. Irradiated discs mutant for the apical caspase Dronc staining with TREred and phalloidin (a), EdU (b) or β-galactosidase activity (c). Note that all TREred cells in the discs are bigger than surrounding cells (a), do not proliferate (b) and show high levels of β-galactosidase activity (c), both in the thoracic and the appendage region. d) Irradiated disc of the genotype *hh^Gal80^>UAS-miRHG, TREred* showing that ROS production, visualized by the expression of the reporter gene *GstD*, takes place in all the TREred cells. e) Phalloidin staining in irradiated disc of the genotype *pnr>UAS-miRHG, TREred* shows that TREred cells generated in the notum region are bigger than surrounding cells. f) EdU incorporation (blue) shows that TREred cells do not divide, in contrast to surrounding cells. g) TREred cells in the notum region produces high levels of β-galactosidase activity assay.

